# Skeletal muscle releases extracellular vesicles with distinct protein and miRNA signatures that accumulate and function within the muscle microenvironment

**DOI:** 10.1101/2021.11.30.470551

**Authors:** Sho Watanabe, Yuri Sudo, Satoshi Kimura, Kenji Tomita, Makoto Noguchi, Hidetoshi Sakurai, Makoto Shimizu, Yu Takahashi, Ryuichiro Sato, Yoshio Yamauchi

## Abstract

Extracellular vesicles (EVs) contain various regulatory molecules and mediate intercellular communications. Although EVs are secreted from various cell types, including skeletal muscle cells, and present in the blood, their identity is poorly characterized *in vivo*, limiting the identification of their origin in the blood. Since the skeletal muscle is the largest organ in the body, it could substantially contribute to circulating EVs as their source. However, due to the lack of defined markers that distinguish SkM-EVs from others, whether the skeletal muscle releases EVs *in vivo* and how much the skeletal muscle-derived EVs (SkM-EVs) account for plasma EVs remain poorly understood. In this work, we perform quantitative proteomic analyses on EVs released from C2C12 cells and human iPS cell-derived myocytes and identify potential marker proteins that mark SkM-EVs. These markers we identified apply to *in vivo* tracking of SkM-EVs. The results show that skeletal muscle makes only a subtle contribution to plasma EVs as their source in both control and exercise conditions in mice. On the other hand, we demonstrate that SkM-EVs are concentrated in the skeletal muscle interstitium. Furthermore, we show that interstitium EVs are highly enriched with the muscle-specific miRNAs and repress the expression of the paired box transcription factor *Pax7*, a master regulator for myogenesis. Taken together, our findings reveal that the skeletal muscle releases exosome-like small EVs with distinct protein and miRNA profiles *in vivo* and that SkM-EVs mainly play a role within the muscle microenvironment where they accumulate.

## Introduction

The skeletal muscle is the largest organ in the body, accounting for 40% of body weight and is responsible for locomotion activity, whole-body metabolism, and energy homeostasis. Moreover, the skeletal muscle serves as a secretory organ (1, 2): it secretes various humoral factors known as myokines, including irisin, apelin, interleukins, and myostatin. They act as mediators for cell-cell communications in autocrine, paracrine, and endocrine fashions. Each myokine has distinct functions and influences tissue homeostasis and metabolism within the skeletal muscle and in other tissues (1, 2). Exercise can induce the expression and secretion of some myokines, which partly explains the health benefits of exercise (1, 3). Thus, the skeletal muscle is considered as an important secretory organ that governs whole-body homeostasis.

In addition to humoral factors, cells release membrane vesicles to the extracellular milieu. Over the last decade, much attention has been paid to the extracellular vesicles (EVs) because they accommodate a wide variety of bioactive molecules, including nucleic acids (DNA, mRNA, microRNA (miRNA), long noncoding RNA), proteins, lipids, and metabolites, and deliver them to recipient cells (4-8). Thus, EVs also act as a means for intercellular and interorgan communications in physiological and pathophysiological settings, including exercise, cancer, and metabolic diseases (9, 10). EVs are heterogeneous in nature and classified into three classes based on size and biogenesis mechanisms, exosomes (50–150 nm in diameter), microvesicles (100– 1,000 nm), and apoptotic bodies (100–5,000 nm) (4-6). Exosomes are derived from the multivesicular bodies (MVBs) of the late endosome. The MVBs fuse with the plasma membrane (PM) and intraluminal vesicles (ILVs) inside the MVBs are released to the extracellular environment as exosomes. Microvesicles are originated from the plasma membrane by membrane budding. Apoptotic bodies are released from apoptotic cells.

EVs are abundantly present in body fluids, including plasma. It is thus expected that their constituents serve as useful biomarkers for diagnosis (9, 11). On the other hand, once they are released from original tissues and enter the circulation, it is nearly impossible to identify their origin because tissue-specific EV markers are poorly characterized. This issue makes it difficult to understand the contribution of each tissue to circulating EVs and to track certain EVs *in vivo*.

Like other cell types, skeletal muscle cells are capable of releasing EVs (12). Evidence shows that C2C12 murine myoblasts and myotubes, and human primary myocytes secret EVs (13-15). EVs released from C2C12 myotubes are transferred to myoblasts and regulate differentiation into myotubes by modulating gene expression (15). Furthermore, EVs derived from C2C12 myotubes contain miRNAs specifically or abundantly expressed in the skeletal muscle called myomiRs (16, 17) that regulate skeletal muscle homeostasis (18, 19). These results suggest that SkM-EVs have physiological functions. Proteomic approaches identified several muscle-specific proteins in C2C12-derived EVs (14, 15). However, their potential as protein markers for SkM-EVs *in vivo* has been poorly explored. Due to the lack of defined SkM-EV markers, whether the skeletal muscle actively releases EVs *in vivo*, how much proportion of plasma EVs are derived from this tissue, and where SkM-EVs are delivered and exert their roles remain largely unknown. In addition, although previous works show that exercise increases circulating EVs (20-22), it is under debate whether skeletal muscle contributes to the exercise-dependent increase in circulating EVs.

To address these issues, here we seek to identify SkM-EV marker proteins by quantitative proteomics on human and mouse myocyte-derived EVs and investigate whether skeletal muscle releases exosome-like small EVs *in vivo*. Based on our proteomic profiling of EVs released from these myocytes, we provide *in vivo* evidence that skeletal muscle actively releases small EVs with distinct protein and miRNA profiles and that SkM-EVs highly accumulate within the skeletal muscle interstitium rather than being secreted into the blood. We further show that EVs isolated from the muscle interstitium modulate myogenic gene expression in murine myoblasts. We thus propose that SkM-EVs mainly exert their functions within the muscle microenvironment.

## Results

### C2C12 cells and hiPSC-derived myocytes secrete EVs

To characterize EVs secreted from both human and mouse skeletal muscle cells, we first isolated EVs from mouse C2C12 myoblasts and myotubes, and human iPSC-derived myocytes (hiPSC-myocytes) by a standard ultracentrifugation protocol (23).

C2C12 myoblasts were differentiated into myotubes (**Figure S1A**) and incubated for 48 h in a differentiation medium containing EV-free horse serum (HS) before isolating EVs from the conditioned medium. To isolate C2C12 myoblast EVs, the cells were incubated for 48 h in an EV-free FBS medium. In addition, we used two lines of hiPSC-myocytes, 414C2^tet-MyoD^ and 409B2^tet-MyoD^. These hiPSC lines harbor tetracycline-inducible human *MYOD1* expressing *piggyBac* vector, and thus adding doxycycline (Dox) into culture medium induces MyoD1 expression, initiating myogenic differentiation. Five–six days after Dox addition, these hiPSCs differentiated into myocytes expressing skeletal muscle cell marker proteins, including myosin heavy chain (MyHC), myogenin, and caveolin-3 but no longer expressing the iPSC marker proteins Nanog, OCT-4A, and Sox 2 (**Figure S1B**). After differentiation, hiPSC-myocytes were incubated in a medium supplemented with EV-free HS for 48 h to isolate EVs from a conditioned medium. To observe the morphology of isolated EVs, we first performed transmission electron microscopy (TEM) analysis on EVs from C2C12 myoblasts (C2C12-MB-EVs), C2C12 myotubes (C2C12-MT-EVs), and hiPSC-myocytes (hiPS-MC-EVs). **Figures 1A** shows typical images of C2C12-MB-EVs, C2C12-MT-EVs, and hiPS-MC-EVs. The diameters of C2C12-MB-EVs and C2C12-MT-EVs were 48.8 ± 16.4 nm and 61.4 ± 22.4 nm, respectively (**Figure 1B**). C2C12-MT-EVs were statistically larger than C2C12-MB-EVs. The average size of hiPS-MC-EVs was approximately 58 nm in both 414C2^tet-MyoD^ and 409B2^tet-MyoD^ lines (**Figure 1B**). The sizes are all within the range of typical exosomes. Together, these results showed that both human and mouse skeletal muscle cells release small EVs with similar morphology.

**Figure 1.**
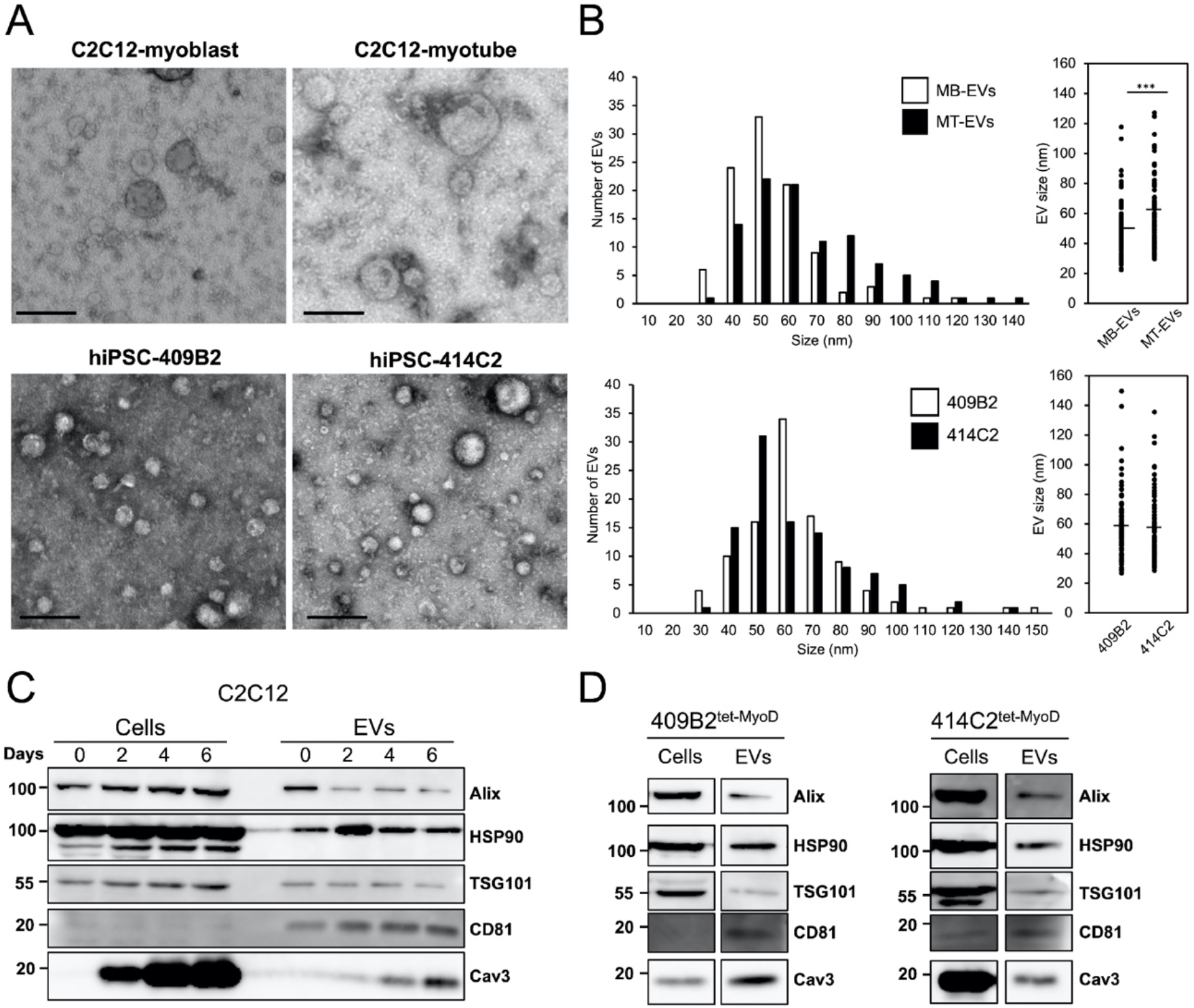
Isolation and characterization of EVs from cultured human and mouse myocytes. (**A**) TEM images of EVs. EVs were isolated from C2C12 myoblasts, C2C12 myotubes, and two lines of hiPSC-myocytes (409B2^tet-MyoD^ and 414C2^tet-MyoD^) and images were acquired under TEM. Scale bar, 100 nm. (**B**) Size distribution of EVs. Sizes of EVs from C2C12-MB-EVs, C2C12-MT-EVs (top), and hiPS-MC-EVs (bottom) were measured using TEM images. Statistical analysis was performed by Student’s *t*-test. ***, *P* < 0.005 (n = 100). (**C**) Differentiation-dependent expression of proteins in C2C12 cells and C2C12-derived EVs. Cell lysates and EVs were prepared on days 0, 2, 4, 6. Forty-eight hours before harvest, medium was switched to EV-free medium. EVs were isolated from conditioned medium by ultracentrifugation as described in the Materials and Methods. Expression of EV marker proteins in cell lysate (20 μg protein) and EVs (1 μg protein) was analyzed by immunoblot. (**D**) Protein expression in hiPSC-myocytes and their EVs. hiPSCs (409B2^tet-MyoD^ and 414C2^tet-MyoD^) were differentiated into myocytes. Afterward, myocytes were incubated for 48 h in medium containing 5% EV-free HS. EVs were isolated from conditioned medium by ultracentrifugation. Cell lysate (20 μg protein) and EVs (1 μg protein) were subjected to immunoblotting to analyze the expression of the indicated proteins.

We next examined the presence of the exosome marker proteins in the isolated EVs. The results show that C2C12-MB-EVs and C2C12-MT-EVs contained the well-defined exosome markers, including Alix, TSG101, CD81, and HSP90 (**Figure 1C**). hiPS-MC-EVs also contained these exosome markers (**Figure 1D**). In addition to these typical markers, we found that C2C12-MT-EVs and hiPS-MC-EVs but not C2C12-MB-EVs, contained caveolin-3, a protein highly expressed in the skeletal muscle and cardiomyocytes, and its contents increased by differentiation. The results suggest that skeletal muscle cells release EVs harboring skeletal muscle-specific proteins.

### Proteomic profiling of EVs released from skeletal muscle cells

To determine proteomic profiling of EVs released by skeletal muscle cells, we first performed quantitative shotgun proteomic analyses on C2C12-MB-EVs and C2C12-MT-EVs. The analyses identified 957 and 1,006 proteins in C2C12-MB-EVs and C2C12 MT-EVs, respectively, which cover 1,047 different proteins (**Figure 2A**). Previously, Forterre *et al*. identified 455 proteins as those found in EVs secreted from C2C12 myoblasts and myotubes (15). Of the 455 proteins, 354 proteins (78%) were also found in our results (**Figure S2**). The current results thus revealed 693 additional C2C12-MB/MT-EVs proteins not identified previously.

**Figure 2.**
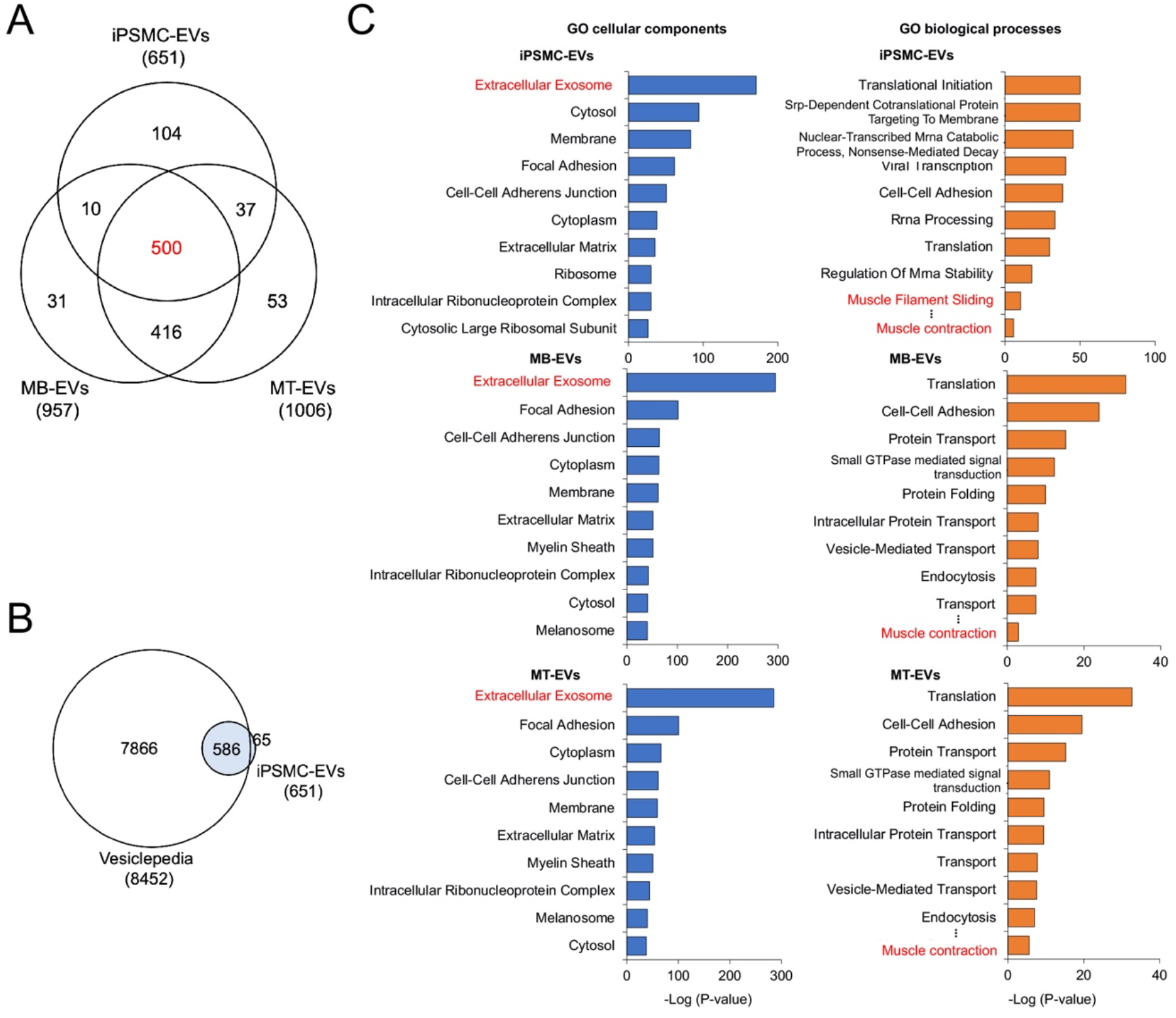
Proteomic profiling of myocyte-derived EVs. (**A**) Venn diagram showing the distinct and overlapping EV proteins from C2C12 myoblasts, C2C12 myotubes, and hiPSC-myocytes. Proteomic analyses were performed on EVs isolated from C2C12 myoblasts (MB-EVs), C2C12 myotubes (MT-EVs), and hiPSC-myocytes (iPSMC-EVs). The 651 proteins in the iPSMC-EVs are derived from EVs isolated from both 414C2^tet-Myo-D^ or 409B2^tet-MyoD^. (**B**) Venn diagram showing proteomic coverage of hiPS-MC-EVs versus Vesiclopedia database. (**C**) GO analysis of myocyte-derived EVs for cellular components (*left*) and biological processes (*right*). Proteomic data on iPS-MC-EVs (*top*), C2C12-MB-EVs (*middle*), and C2C12-MT-EVs (*bottom*) were analyzed using DAVID. Top 10 GO term are listed.

Next, the same proteomic analysis was performed on hiPS-MC-EVs from both 414C2^tet-Myo-D^ and 409B2^tet-MyoD^ lines, and 651 proteins were detected (**Figure 2A**). Among these 651 proteins, the 586 proteins (90%) are covered by the EV database Vesiclepedia (24) (http://microvesicles.org), validating isolated EVs of quality (**Figure 2A**). Five hundred forty-seven proteins (84%) out of the 651 were also found in EVs isolated from either C2C12-MB-EVs or C2C12-MT-EVs (**Figure 2A**). The results also identified 500 proteins that overlap among C2C12-MB-EVs, C2C12-MT-EVs, and hiPS-MC-EVs. Thirty-seven proteins (**Table S1**) were found in both C2C12-MT-EVs and hiPSM-EVs but not in C2C12-MB-EVs, suggesting a distinct protein profile of myotube-derived EVs. On the other hand, 31 proteins (**Table S2**) were found only in C2C12-MB-EVs.

To annotate identified EV proteins, we classified these proteins based on Gene Ontology (GO) using an integrative platform, DAVID (25, 26). The results showed that in hiPS-MC-EVs, C2C12-MB-EVs, and C2C12-MT-EVs, proteins belonging to the term “Extracellular Exosome” in “Cellular Components” were highly enriched, confirming that isolated EVs are of good quality (**Figure 2C**). For the “Biological processes” term, proteins classified into “Muscle contraction” were significantly enriched in all three EV samples, which indicates that SkM-EVs contain proteins unique to the skeletal muscle. Together, all these results suggest that SkM-EVs display a distinct protein signature.

### Identification of potential marker proteins for SkM-EVs

We next sought to identify potential marker proteins that mark EVs released from skeletal muscle cells. To this end, we searched proteins highly expressed in the skeletal muscle from our proteome data obtained from hiPS-MC-EVs and C2C12-MT-EVs. As mentioned above, we identified thirty-seven potential MT-EV proteins (**Figure 2A, Table S1**). To assess their specificity, we searched specific proteins using the Gene Ontology Consortium’s Community Annotation Wiki for Muscle Biology (http://wiki.geneontology.org/index.php/Muscle_Biology) and confirmed that many of these proteins, including Nebulin, KLHL41, MYH1, TRIM72, ACTA1, and MYBPH are predominantly expressed in the skeletal muscle. Furthermore, based on The Human Protein Atlas and The Genotype-Tissue Expression (GTEx) databases, we selected 10 proteins as potential marker proteins for SkM-EVs (**Figure 3A**). To confirm whether these proteins are included in EVs, C2C12-MT-EVs were isolated by the previously defined Tim4-based method (27) and subjected to immunoblot analysis. Due to the availability and/or validity of antibodies, six out of ten proteins were analyzed. The results show that in addition to the typical exosome marker proteins (Alix, Annexin A1, CD81, and Flotillin-1), C2C12-MT-EVs contain the skeletal muscle proteins, ATP2A1, β-enolase, calsequestrin 2, caveolin-3, and desmin (**Figure 3B**), validating our proteomic analysis. Among them, ATP2A1, β-enolase, and desmin are predominantly expressed in skeletal muscle tissues (**Figure S3**).

**Figure 3.**
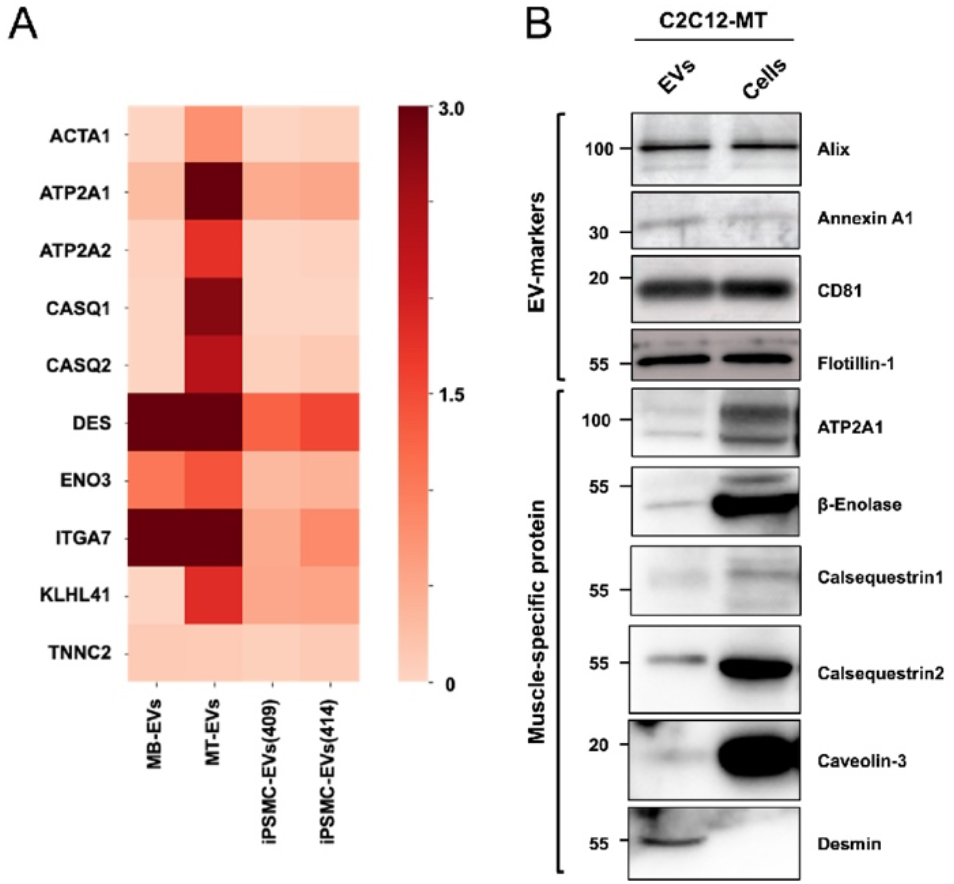
Identification of potential SkM-EV markers *in vitro*. (**A**) Heatmap showing the contents of potential SkM-EV marker in isolated EVs. The contents of the ten muscle-specific proteins in C2C12-MB-EVs (MB-EVs), and C2C12-MT-EVs (MT-EVs), and hiPS-MC-EVs (iPSMC409-EVs and iPSMC414-EVs) are shown. (**B**) Expression of EV-marker proteins in C2C12-MT-EVs. EVs and cell lysate were prepared using PS-affinity beads and urea buffer, respectively. The expressions of typical EV marker proteins and muscle-specific proteins in EVs and cells were analyzed by immunoblot.

### SkM-EVs accumulate in the skeletal muscle interstitium

Recent reports showed that apart from plasma, the interstitium of tissues such as the liver and lung contain significant amounts of EVs (28). To determine whether the skeletal muscle cells release EVs *in vivo*, we isolated EVs from both plasma and skeletal muscle (tibialis anterior, gastrocnemius, soleus, and quadriceps) interstitium of mice using the Tim4-based method (**Figure 4A**). We validated the quality of isolated EVs by TEM and confirmed that EVs from both the plasma and skeletal muscle interstitium show similar morphology (**Figure 4B**). Plasma and SkM-interstitium EVs were similar in size, ranging from 30–150 nm (**Figure 4C**). Scanning electron microscopy (SEM) analysis showed that the skeletal muscle interstitium contains EV-like vesicles with a diameter of 50–500 nm, which are attached to extracellular matrix (ECM)-like structures (**Figure 4D**). We next examined whether these EVs contain SkM-EV markers identified above. As expected, the typical EV marker protein Alix and CD81 were found in both plasma and SkM-interstitium EVs (**Figure 4E**). In addition, SkM-EV marker proteins were detected at high levels in the SkM-interstitium EVs. In contrast, the SkM-EV markers were much less or undetectable in the plasma EVs. ATP2A1 and desmin were slightly detected in plasma EVs, suggesting that SkM-EVs only partly enter the bloodstream. These results indicate that SkM-EVs are highly concentrated in skeletal muscle tissues but are not major populations in the circulation.

**Figure 4.**
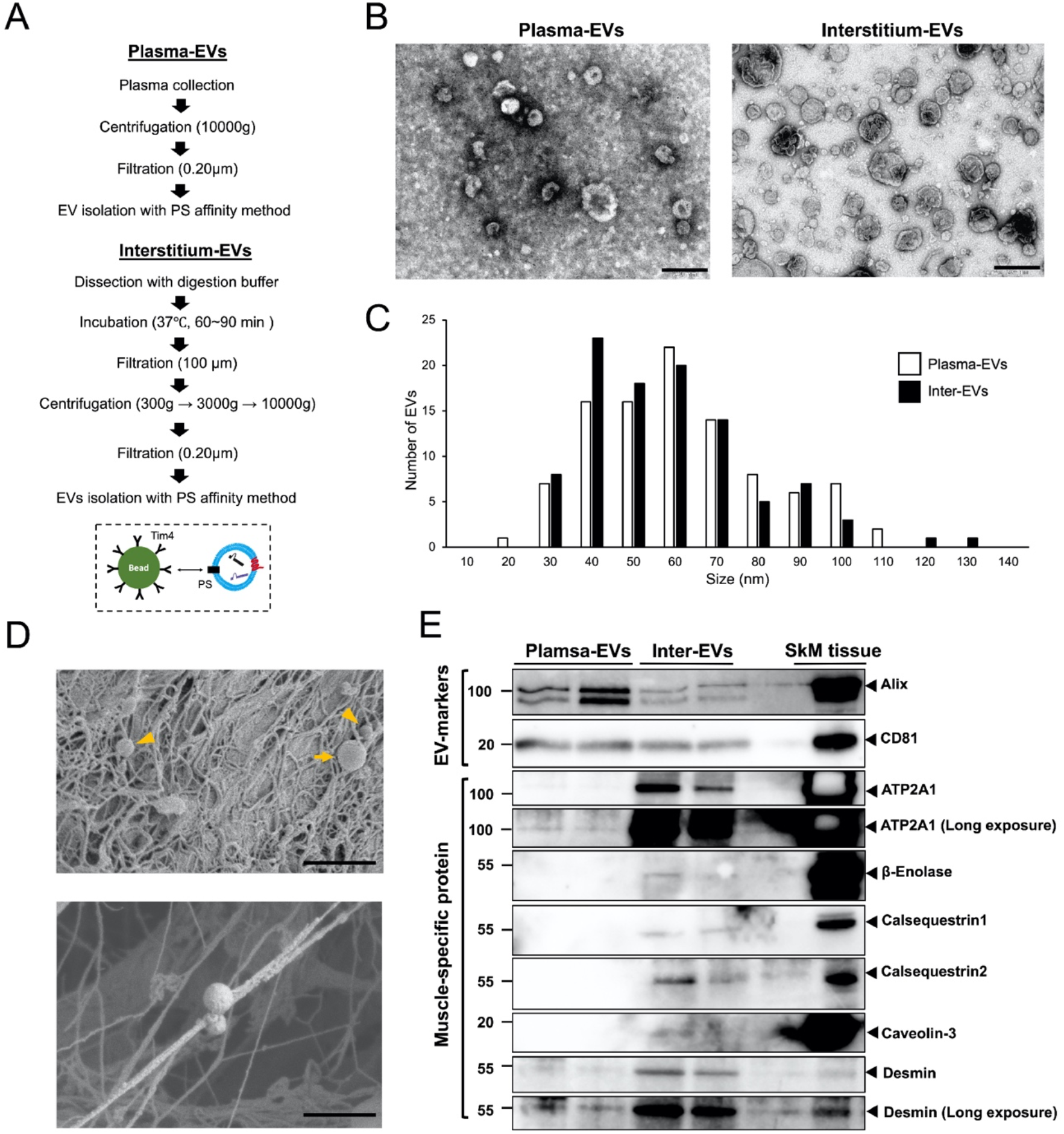
Validation of SkM-EVs markers *in vivo*. (**A**) Outlines of EV isolation protocols from plasma and the skeletal muscle interstitium. See the Material and Methods for more detail. (**B**) TEM images of plasma and interstitium EVs. Scale bar, 200 nm. (**C**)Size distribution of plasma and interstitium EVs. Size of EVs were analyzed using Image J. (**D**) SEM images of skeletal muscle tissue (gastrocnemius). Small and large EVs are indicated arrowheads and arrows, respectively. The bottom image shows small EVs attaching ECM-like structures. Scale bar, 1 μm in upper panel and 500 nm in lower panel. (**E**) Expression of the EV marker proteins in plasma and interstitium EVs. Plasma and interstitium EVs were isolated from two mice as in (A). Skeletal muscle tissue (quadriceps) homogenates were also prepared from the same mice. Plasma EVs (5 μg protein/lane) and interstitium EVs (Inter-EVs) (5 μg protein/lane) were subjected to immunoblot analysis to validate the presence of the marker proteins in these EVs. SkM tissue homogenates (2 μg protein/lane) were also analyzed as positive controls.

To further determine the physiological importance of SkM-EVs *in vivo*, we examined the effect of exercise on SkM-EVs. Whether exercise increases circulating EV contents is currently controversial (29-31). Moreover, even though exercise increases circulating EVs, their origin(s) is not fully characterized. We took advantage of our newly identified SkM-EV marker proteins to clarify this issue. After mice were subjected to exhaustive endurance running on a treadmill (**Figure S4A, B**), we immediately harvested blood from the heart and skeletal muscle tissues from a hind limb, and prepared plasma EVs and SkM-interstitium EVs, respectively. The results showed that exercise does not alter protein contents in either plasma EVs (Ctrl: 340.3 ± 22.5 mg/mL; Exercise: 328.4 ± 8.0 mg/mL; *p* = 0.30) or SkM-interstitium EVs (Ctrl: 371.4 ± 19.5 mg/mL; Exercise: 378.7 ± 23.6 mg/mL; *p* = 0.35). We also assessed levels of marker proteins in plasma and the interstitium EVs. Neither typical EV markers nor SkM-EV markers were not changed by exercise in plasma and the interstitium EVs (**Figure 5A, B**). Exercise did not influence the expression of SkM-EV marker proteins in the skeletal muscle (**Figure S4C**). These results suggest that exercise does not influence EV release from the skeletal muscle. Meanwhile, we observed positive correlations between CD81 and β-enolase, caveolin-3, or ATP2A1 contained in the interstitium EVs (**Figure 5C, Table S3**). It could be consistent with previous studies that skeletal muscle cells preferentially release CD81-positive EVs (12, 13). Furthermore, we noticed subtle but significant increases in the size of the interstitium EVs but not of plasma EVs upon exercise (**Figure 5D, E**). Together, our results show that the skeletal muscle releases EVs with a distinct protein signature and that SkM-EVs highly accumulate in the interstitium. Our results also reveal that SkM-EVs do not account for the major proportion of circulating EVs.

**Figure 5.**
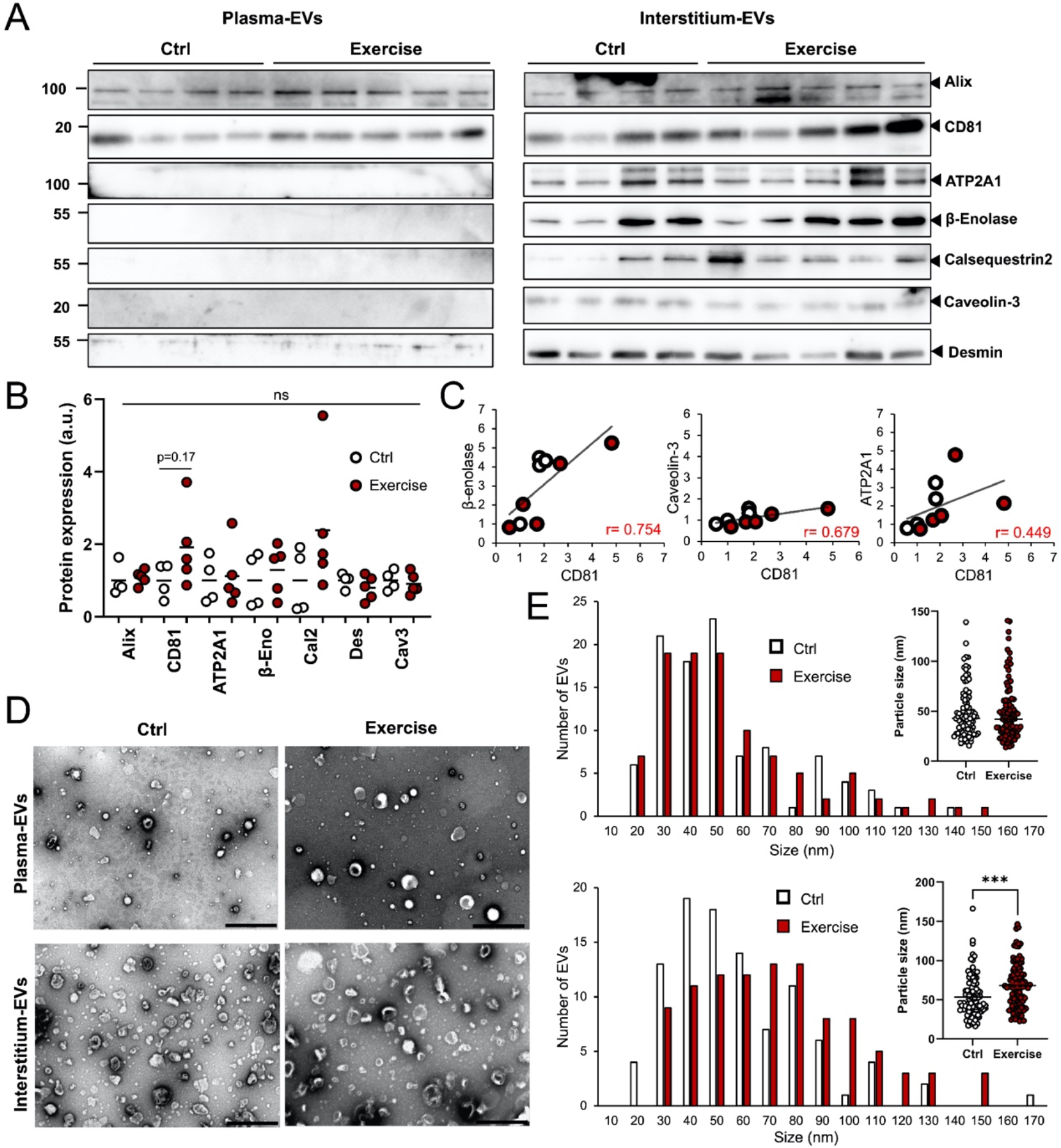
Effect of exercise on plasma and SkM interstitium EVs. (**A**) Immunoblot analysis of EVs. Plasma and skeletal muscle interstitium EVs were isolated from control (n = 4) and exercised mice (n = 5). Equal volume of plasma EVs (20 μL/lane, equivalent to EVs from 60 μL plasma) and interstitium EVs (20 μL/lane, equivalent to EVs from 60 mg tissue) were subjected to immunoblotting to detect the indicated proteins. Expression of these proteins in skeletal muscle tissues with or without exercise are shown in Figure S4C. (**B**) Quantification of protein levels in interstitium EVs before and after exercise. (**C**) Correlation between CD81 and SkM-EV marker proteins (β-enolase, caveolin-3, and ATP2A1). Each circle represents individual mice with (red circle) or without (white circle) exercise. (**D**) TEM images of plasma and interstitium EVs with or without exercise. Scale bar, 500 nm. (**E**) Size distribution of plasma (*upper*) and interstitium (*lower*) EVs isolated from mice with or without exercise. Statistical analysis was performed by Student’s *t*-test. ***, *P* < 0.005 (n = 100).

### SkM-interstitium EVs are rich in myomiRs and promote myoblast differentiation

EVs are characterized as the vehicle for miRNAs. Therefore, we finally investigated myomiR profiles of SkM-interstitium EVs and plasma EVs. The results show that all the four miRNAs (miRs-1, -206, -431, and -486) abundantly expressed in the muscle are markedly concentrated in the interstitium EVs (**Figure 6A**). In particular, miR-1 and miR-206 in the interstitium EVs were 45- and 20-fold higher than those in plasma EVs, respectively, confirmaing the intramuscular accumulation of SkM-EV detected by our protein-based analysis. Together, these results demonstrate that SkM-interstitium EVs display unique protein and miRNA profiles that are distinct from plasma EVs. MyomiRs play important roles in skeletal muscle homeostasis, including the regulation of myogenesis by targeting the paired box transcription factor Pax7, a master regulator for myogenesis (32-34). Our results led us to hypothesize that SkM-EVs predominantly play their roles within the intramuscular microenvironment. To test this, we asked whether SkM-interstitium EVs isolated from mice modulate the expression of genes involved in myogenesis. **Figure 6B** shows that C2C12 myoblasts uptake the interstitium EVs, suggesting that SkM-EVs function in these cells. We next determined mRNA levels that regulate myoblast differentiation. The results show that the interstitium EVs suppress *Pax7* expression but increase *MyHC* expression, a marker for myoblast differentiation (**Figure 6C**). Although the downregulation of *Pax7* by interstitium EVs was not as robust as overexpression of myomiRs in myoblasts, our results were largely consistent with previous studies showing the repression of *Pax7* mRNA expression by myomiRs (32-34). The current results thus suggest that SkM-interstitium EVs regulate myogenesis at least in part by suppressing *Pax7* expression within the muscle microenvironment (**Figure 6D**).

**Figure 6.**
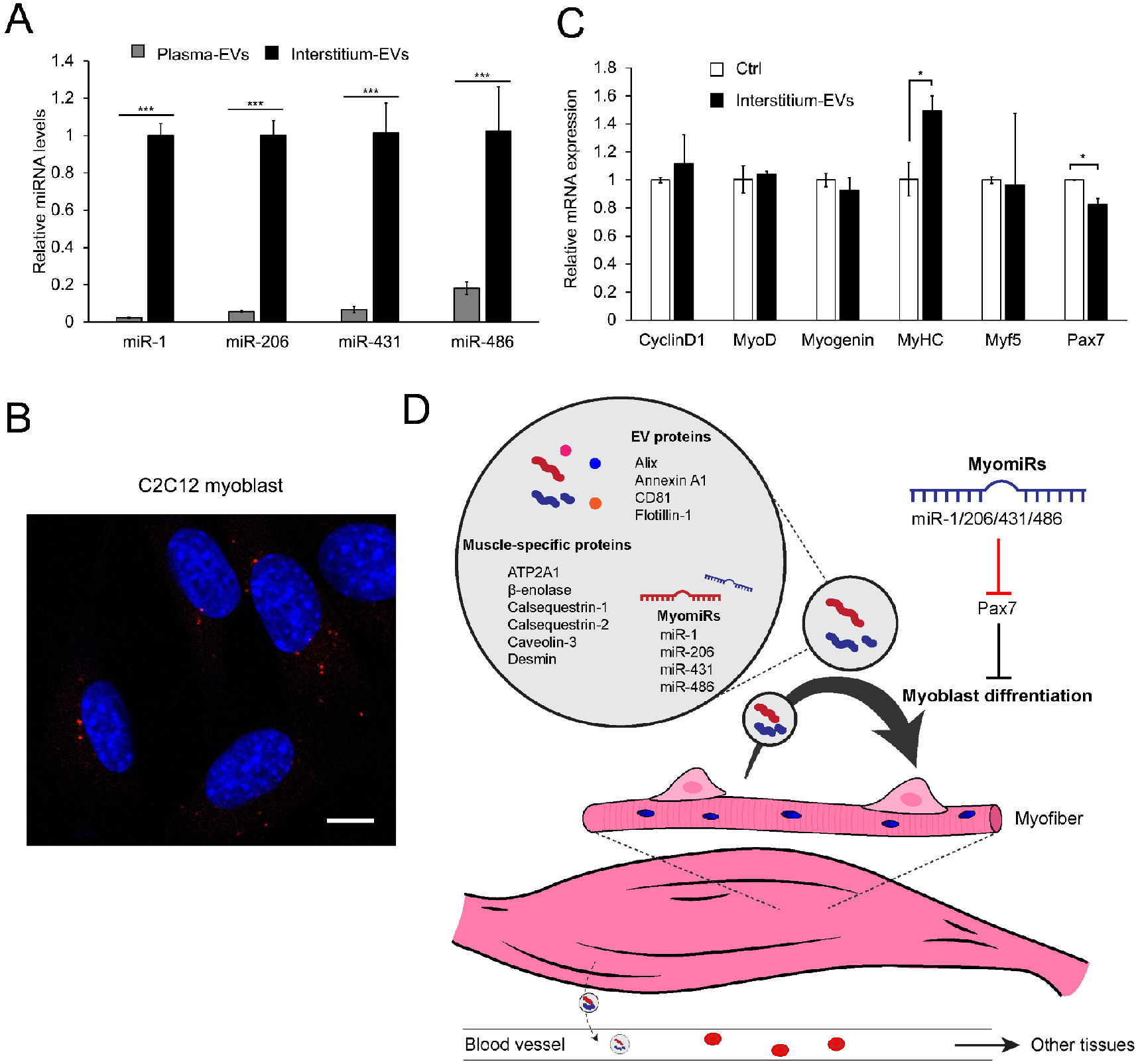
EVs uptake and functional analysis of SkM-interstitium EVs. (**A**) Exosomal miRNAs. Levels of miRs-1, -206, -431, and -486 in plasma and interstitium EVs were determined by qPCR as described in Materials and Methods. (**B**) Uptake of interstitium EVs by myoblasts. C2C12 myoblasts were incubated with labeled-interstitium EVs (4 μg protein/well, red) for 6 h. After fixation, permeabilization, and nucleus staining with DAPI (blue), cell images were acquired by a confocal microscopy. Bar, 10 μm. (**C**) Effect of interstitium EVs on gene expression in myoblasts. C2C12 myoblasts were incubated with interstitium EVs (4 μg protein/well) in growth medium for 24 h. mRNA levels of the indicated genes were analyzed by qPCR. Results are shown as mean ± SEM (*n* = 3). Statistical analysis was performed by Student’s *t*-test. * *P* < 0.05, ** *P* < 0.01, *** *P* < 0.01. (**D**) A model depicting the role of SkM-EVs. See text for more detail.

## Discussion

Circulating EVs are derived from various sources. For identification of their origin, it is essential to determine tissue-specific EV marker molecules. Proteomic approaches have been taken to search tissue-specific EV markers using cell culture models, including skeletal muscle cells (13, 15), hepatocytes (35), and adipocytes (36). Although these studies identified potential tissue-specific EV marker proteins, their validity *in vivo* has been poorly characterized. Accordingly, much less is known about the dynamic movement of EVs in the body. In this work, we sought to identify marker proteins that help characterize EVs derived from the skeletal muscle both in humans and mice. We first determined proteomic profiles of EVs released from C2C12 myoblasts, C2C12 myotubes, and hiPSC-myocytes and identified several proteins that serve as potential markers for SkM-EVs. We then demonstrated that these marker proteins are relevant to identifying SkM-EVs *in vivo*. Finally, we showed that SkM-EVs accumulate within the muscle microenvironment where they regulate gene expression, rather than enter the blood circulation.

In addition to myokines secreted upon exercise, exercise-induced EVs are expected to exert health benefits (37, 38). A recent work showed that SkM-EVs from trained mice contain higher levels of miR-133b, which suppresses FoxO1 expression in the liver and improves insulin sensitivity (28). On the other hand, it was shown that the skeletal muscle is not the major source of exercise-induced EVs (22). It is thus important to identify the nature of exercise-induced EVs, including their components and origins.

Attempts have been made to identify markers for SkM-EVs, yet any defined markers applicable to *in vivo* analysis have not been determined at present. Several studies have reported potential markers for SkM-EVs. It was suggested that α-sarcoglycan (SGCA)-positive EVs present in the plasma are derived from the skeletal muscle (39). In contrast, other studies failed to detect SGCA-positive EVs in human subjects either before or after exercise training (22). Furthermore, SGCA is not exclusively expressed in the skeletal muscle but also expressed in other tissues, including the heart, smooth muscle, and lung. In addition, although myomiRs are found in both human and mouse plasma EVs (16), a recent work demonstrates that adipose tissue is a major source of circulating exosomal miRNAs (40). These reports suggest that skeletal muscle makes only a subtle contribution to circulating EV and miRNA levels quantitatively. Meanwhile, these contrasting findings indicate the lack of consensus on how SkM-EVs behave after secretion and the difficulty in tracking SkM-EVs *in vivo*.

Our current analyses on proteomic profiling of human and mouse skeletal muscle cell-derived EVs combined with *in vivo* validation provide more reliable markers for SkM-EVs. Among proteins predominantly expressed in the skeletal muscle, we showed that ATP2A1, β-enolase, and desmin may serve as reliable SkM-EV marker proteins. By monitoring these marker proteins, we investigated whether SkM-EVs account for a significant proportion of circulating EVs and whether exercise increases SkM-EVs *in vivo*. Unexpectedly, SkM-EVs marker proteins were hardly detected in the plasma even after exercise. Consistent with this observation, our exosomal miRNA analysis showed that myomiR levels in plasma EVs are only subtle compared to those in interstitium EVs. These results could be consistent with previous reports showing that SGCA-positive EVs constitute only 1–5 % of total circulating EVs (39) and that most circulating exosomal miRNA are derived from adipose tissue (40). Our results are also supported by evidence that exercise-induced EVs are derived from leukocytes, platelets, and endothelial cells (22) and that treadmill running does not influence muscle-specific miRNA levels in serum (41). In contrast, we found that SkM-EVs are highly accumulated in the skeletal muscle interstitium. All these results support our view that the skeletal muscle is not the major source of circulating EVs regardless of physical activities and that SkM-EVs dominantly play a role within the tissue, not at systemic levels (**Figure 6D**).

What is the role of SkM-EVs in the muscle microenvironment? Our data disclosed that SkM-interstitium EVs contain myomiRs at very high levels. The myomiRs we found in the interstitium EVs serve as negative regulators of Pax7, leading to myoblast differentiation (32-34). We showed that SkM-interstitium EVs isolated from mice suppress *Pax7* gene expression and up-regulate *MyHC* gene expression in murine myoblasts. We thus propose that SkM-EVs support myogenesis at least partly through myomiRs-mediated suppression of Pax7 expression. Although our data showed that only subtle amounts of SkM-EVs are found in the blood, it was reported that SkM-interstitium EVs modulate hepatic gene expression when added in cultured hepatocytes or injected intravenously in mice (28). Whether sufficient amounts of SkM-EVs are delivered to other tissues through the circulation for regulating the physiological states of recipient tissues/cells may need further investigation.

In summary, we revealed the distinct protein and miRNA profiles of SkM-EVs *in vivo*. Tracking SkM-EV markers led us to conclude that SkM-EVs do not account for the major population of circulating EVs although the skeletal muscle is the largest tissue in the body. Rather, we showed that SkM-EVs highly accumulate within the skeletal muscle microenvironment where they regulate gene expression to promote myogenesis at least partially through myomiRs.

## Materials and Methods

### Materials

Fetal bovine serum (FBS) and horse serum (HS) were obtained from Gibco. FBS and HS were heat-inactivated before use. EV-free FBS and HS were prepared as described (23). Briefly, FBS and HS were spun at 2,000 *g* for 10 min followed by centrifugation at 100,000 *g* for 70 min. The supernatant was further centrifuged at 100,000 *g* for 16 h. The supernatant was filtrated with a 0.20 μm filter (Advantec) and used as EV-free FBS or HS. EV-free FBS and HS were stored at -80 °C until use.

### Cell culture

C2C12 mouse myoblasts (obtained from ATCC) were maintained at low cell density in growth medium (DMEM supplemented with 10% FBS). For differentiation to myotubes, C2C12 myoblasts were seeded into a 6-well plate at a density of 1.5 × 10^5^ cells per well and grown for 2 days in a growth medium. Afterward, cells were incubated for 4 days in a differentiation medium (DMEM supplemented with 2% HS). The medium was changed every other day. For isolation of EVs, cells were incubated in 2% EV-free HS for 48 h. Human iPS cell (hiPSC) lines, 414C2^tet-Myo-D^ and 409B2^tet-MyoD^ were maintained in StemFit AK02N (Ajinomoto) as described (42). These hiPSCs were differentiated into myocytes by a published protocol (42). Briefly, on day 0, hiPSCs were seeded into a Matrigel-coated 6-well plate at a density of 3–4 × 10^5^ cells/well and grown overnight in StemFit medium with 10 μM Y-27632. On day 1, the medium was switched to Primate ES Cell Medium (Reprocell) containing 10 μM Y-27632. On day 2, cells were incubated in Primate ES Cell Medium containing 1 μg/mL doxycycline (Dox) to induce MyoD1 expression. On day 3, the medium was changed to αMEM containing 5% KnockOut Serum Replacement (Gibco) and Dox (1 μg/mL) and incubated for 2–3 days. After differentiation, hiPSC-myocytes were incubated in DMEM containing 2% EV-free HS for 48 h to isolate EVs.

### Animal studies

All protocols for animal procedures were approved by the Animal Care and Use Committee of the University of Tokyo, which are based on the Law for the Humane Treatment and Management of Animals (Law No. 105, 1 October 1973, as amended on 1 June 2020). C57BL/6J male mice at 8 weeks old were obtained from Japan Clea. Mice were housed in a 12 h-light/12 h-dark schedule at 23 ± 2°C and 55 ± 10% humidity and fed *ad libitum* with a standard chow diet (Labo MR Stock, Nosan Corporation) and water. Mice at 9–10 weeks old were randomly assigned to either exercise or sedentary groups. After mice were adapted to the treadmill (5 m/min for 10 min per day) for 4 days, they were subjected to exhaustion running for up to 90 min using a ramped treadmill exercise protocol starting at 10 m/min and increasing by 2 m/min every 10 min (21) using a treadmill (MK-680C, Muromachi Kikai). Mice were defined as the exhausted state when they stopped running on a treadmill for more than 5 s despite gentle encouragement.

Immediately after exercise, blood was collected by cardiac puncture under anesthesia with isoflurane. Afterward, mice were perfused through the left ventricle with PBS for 2 min at a rate of 1 mL/min to remove blood from the tissue, and skeletal muscle (tibialis anterior, gastrocnemius, soleus, quadriceps) and other tissues were then harvested.

### Isolation of EVs from conditioned media

We used two methods to isolate EVs. Method I: EVs were isolated by ultracentrifugation according to a method previously described (23). Briefly, conditioned media (typically 12 mL from 6 wells) where cells were incubated in EV-free medium for 48 h was spun sequentially at 300 *g* for 10 min, 2,000 *g* for 10 min and 10,000 *g* for 30 min. After each centrifugation step, the supernatant was transferred to a new centrifuge tube. The 10,000 *g*-supernatant was filtered through a 0.20 μm filter (Advantec) to obtain small EVs. Afterward, the supernatant was ultracentrifuged at 100,000 *g* for 70 min at 4°C using an MLA-55 rotor (Beckman Coulter) and an Optima MAX-TL Ultracentrifuge (Beckman Coulter). Pellet was washed once with PBS (2 mL/tube) and EVs were pelleted by ultracentrifugation at 100,000 g for 70 min at 4°C again. Resulting pellet was resuspended in 150 μL of PBS. Method II: The 10,000 *g*-supernatant was prepared as described above. After filtration and concentration with Amicon Ultra-15 (Merck), EVs were isolated using by MagCapture Exosome Isolation Kit PS (Fujifilm-Wako) according to the manufacture instruction. This method is based on the ability of Tim4 protein to bind phosphatidylserine (PS) which localizes on the exosome surface (27). In brief, medium concentrated (1 mL) as above was mixed with 0.6 mg of streptavidin magnetic beads bound to 1 μg of biotinylated mouse Tim 4-Fc and incubated in the presence of 2 mM CaCl_2_ for 16–18 h with rotation at 4 °C. After washing beads three times with 1 mL of washing buffer (20 mM Tris-HCl pH 7.4, 150 mM NaCl, 2 mM CaCl_2_, 0.0005% Tween20), exosomes (EVs) were eluted twice with 50 ml of elution buffer (20 mM Tris-HCl pH 7.4, 150 mM NaCl, 2 mM EDTA). EV protein contents were determined by Micro BCA Protein Assay Kit (Thermo Fisher). EVs were stored at -80 °C until use.

### Isolation of EVs from mice

Plasma EVs were isolated by MagCapture Exosome Isolation Kit as above. Plasma (300 μL) was mixed with PBS (600 μL) and spun at 10,000 *g* for 30 min. After filtration of the supernatant with a 0.20 μm filter, plasma was subjected to the isolation of EVs using MagCapture Exosome Isolation Kit with elution volume of 100 μL per 300 μL plasma. Skeletal muscle interstitium EVs were isolated according to a method recently reported (43, 44). Approximately 150 mg of skeletal muscle tissues (tibialis anterior, gastrocnemius, soleus, and quadriceps) from a hind limb were combined and digested with collagenase (10 mg/mL, Sigma) and dispase II (10,000 PU/mL, Wako-Fujifilm) for 1 h at 37°C in HEPES buffer (100 mM HEPES, 2.5 mM CaCl_2_). To avoid disruption of cells, tissues were minced gently. Afterward, one volume of PBS containing 2 mM EDTA was added to the sample, and the sample was passed through a 100 μm cell strainer (Corning). Samples were then centrifuged at 600 *g* for 5 min at 4°C, 2,000 *g* for 10 min, and 10,000 *g* for 30 min. The supernatant was filtrated with a 0.20 μm filter and concentrated using Amicon Ultra-15 (Merck). EVs were then isolated by MagCapture Exosome Isolation Kit as described above. EVs were eluted with 100 μL of elution buffer per 300 mg tissue.

### Immunoblotting and antibodies

Cells were lysed with urea buffer (8 M Urea, 50 mM Na-phosphate pH 8.0, 10 mM Tris-HCl pH 8.0, 100 mM NaCl) containing protease inhibitor cocktail (Nacalai Tesque) as described (45). Tissue homogenates were prepared in radioimmunoprecipitation assay buffer (50 mM Tris-HCl pH 7.4, 150 mM NaCl, 1 mM EDTA, 1% Nonidet P-40, and 0.25% sodium deoxycholate) supplemented with a protease inhibitor mixture (Nacalai Tesque) and phosphatase inhibitor mixture (Sigma). Protein concentration was determined by BCA Protein Assay (Thermo Fisher). Cell lysate, tissue homogenate, and EVs were mixed with Laemmli buffer and heated at 95 °C for 3 min. Aliquots were subjected to SDS-PAGE and immunoblot analysis according to a standard protocol. The expression of a protein was analyzed by Image J software or Evolution-Capt software (Vilber Lourmat). Antibodies used were obtained from commercial sources as follows: anti-caveolin 3 (sc-5310), anti-calsequestrin 1 (sc-137080), anti-calsequestrin 2 (sc-390999), anti-CD81 (sc-166029), anti-β enolase (sc-100811), anti-HSP90 (sc-13119), anti-tsg101 (sc-7964) antibodies from Santa Cruz Biotechnology; anti-flotillin 1 antibody (ab41927) from Abcam; anti-CD81 (#10037), anti-Alix (#92880), anti-desmin (#5332), anti-ATP2A1 (#12293), anti-GAPDH (#5174), HRP-linked anti-mouse IgG (#7076), and HRP-linked anti-rabbit IgG (#7074) antibodies from Cell Signaling Technology; anti-annexin A1 mouse mAb (66344-1-Ig) from Proteintech; Mouse TrueBlot: Anti-Mouse Ig HRP (18-8817-31) from Rockland Immunochemicals.

### Proteomic analysis of EVs

EVs were solubilized in 50 mM Tris-HCl pH 9.0 containing 5% sodium deoxycholate, reduced with 10 mM dithiothreitol for 60 min at 37 °C, and alkylated with 55 mM iodoacetamide for 30 min in the dark at 25 °C. The reduced and alkylated samples were diluted 10-fold with 50 mM Tris-HCl pH 9.0 and digested with trypsin at 37 °C for 16 h (trypsin-to-protein ratio of 1:20 (w/w)). An equal volume of ethyl acetate was added to each sample solution and the mixtures were acidified with the final concentration of 0.5% trifluoroacetic acid. The mixtures were shaken for 1 min and centrifuged at 15,700 g for 2 min. The aqueous phase was collected and desalted with C18-StageTips. LC-MS/MS analysis was performed using an UltiMate 3000 Nano LC system (Thermo Fisher Scientific) coupled to Orbitrap Fusion Lumos hybrid quadrupole-Orbitrap mass spectrometer (Thermo Fisher Scientific) with a nano-electrospray ionization source. The sample was injected by an autosampler and enriched on a C18 reverse-phase trap column (100 μm × 5 mm length, Thermo Fisher Scientific) at a flow rate of 4 μL/min. The sample was subsequently separated by a C18 reverse-phase column (75 μm × 150 mm length, Nikkyo Technos) at a flow rate of 300 nL/min with a linear gradient from 2% to 35% mobile phase B (95% acetonitrile and 0.1% formic acid). The peptides were ionized using nano-electrospray ionization in positive ion mode. The raw data were analyzed by Mascot Distiller v2.3 (Matrix Science), and peak lists were created based on the recorded fragmentation spectra. Peptides and proteins were identified by Mascot v2.3 (Matrix Science) using UniProt database with a precursor mass tolerance of 10 ppm, a fragment ion mass tolerance of 0.01 Da and strict trypsin specificity allowing for up to 1 missed cleavage. The carbamidomethylation of cysteine and the oxidation of methionine were allowed as variable modification.

### Electron microscopy

Specimens for transmission electron microscopy (TEM) were prepared at room temperature. An aliquot of EV sample was pipetted onto a copper grid with carbon support film and incubated for 10 min. After the excess liquid was removed, a grid was briefly placed on 10 μL 2% uranyl acetate (w/v, Merck). Images were acquired under a JEM-1010 electron microscope (JEOL) operated at 100 kV with a Keen view CCD camera (Olympus Soft Imaging Solution). The size of EVs was measured using Image J software.

For scanning electron microscopy (SEM) analysis, skeletal muscle tissue (approximately 3 × 3 mm in size) was fixed with 10% neutral buffered formalin for 1 h and with 0.2 % glutaraldehyde and 2% paraformaldehyde in PBS for 1 h. After post-fixation with 1 % osmium tetroxide in PBS, samples were dehydrated in ethanol series (70%, 90%, 95%, 99.5% and 100%) for 10 min each, treated with tert-butyl alcohol for 10 min twice and freeze-dried. The dried specimen was applied onto a carbon double side-tape with silver paste and sputter coated with platinum palladium. Images were acquired under a Hitachi S-4800 scanning electron microscope with a secondary electron in-lens detector.

### EV labeling and confocal microscopy

EVs were labeled using ExoSparkler Exosome Protein Labeling Kit-Red (Dojindo Laboratories) according to the manufacturer’s instruction. C2C12 myoblasts seeded in a 35-mm film bottom dish (Matsunami) were incubated without or with the labeled EVs (4 μg protein per 2 mL). Cells were fixed with 4% paraformaldehyde (Fujifilm-Wako) for 10 min and then permeabilized with 0.1% Triton X-100 in PBS for 5 min at room temperature. After nuclei were stained with DAPI, specimens were mounted with ProLong Gold Antifade Reagent (Thermo Fisher). Cell images were acquired by an LSM800 confocal laser microscope (Carl Zeiss) equipped with a Plan-Apochromat 63x/1.4 objective. Images were processed with a Zen software (Carl Zeiss).

### mRNA expression analysis

Total RNA was isolated using ISOGEN (NIPPON GENE), according to the manufacturer’s instructions. The high-capacity cDNA reverse transcription kit (Applied Biosystems) was used to synthesize cDNA from total RNA. Quantitative real-time PCR (qPCR) analyses were performed using an Applied Biosystems StepOnePlus. mRNA levels were normalized to 18S ribosomal RNA levels. The primers used for qPCR analysis are described in *SI Appendix*, Table S4.

### miRNA analysis

Plasma and SkM-interstitium EVs were isolated from two mice and pooled for miRNA analyses. Total RNA was extracted from EVs (20 μg protein) using miRNeasy Mini Kit (Qiagen). RNA was then reverse-transcribed using TaqMan MicroRNA Reverse Transcription Kit (Applied Biosystems) according to the manufacturer’s protocol. qPCR was then performed using TaqMan MicroRNA Assay (miR-1, Assay ID: 002222; miR-206, Assay ID: 000510; miR-431, Assay ID: 001979; miR-486, Assay ID: 002093; miR-16, Assay ID: 000391; miR-21, Assay ID: 000397) (Applied Biosystems). Exosomal miRNA levels were normalized by the mean value of miR-16 and miR-21 as described (28).

### Statistical analysis

Results are expressed as mean ± SEM from at least three independent biological replicates. Statistical analyses were performed using the two-tailed, unpaired Student’s *t*-test. *P* values less than 0.05 were considered statistically significant.

## Acknowledgments

This work was supported by KAKENHI grants 19H02908 (to Y.Y.) and 20H00408 (to R.S.) from the Japan Society for the Promotion of Science, and AMED-CREST grants 20gm091008h and 21gm091008h (to Y.Y. and R.S.) from Japan Agency for Medical Research and Development. S.W. was supported by Japan Society for the Promotion of Science Research Fellowship for Young Scientists.

