## Supporting Information for "Skeletal muscle releases extracellular vesicles with distinct protein and miRNA signatures that accumulate and function within the muscle microenvironment"

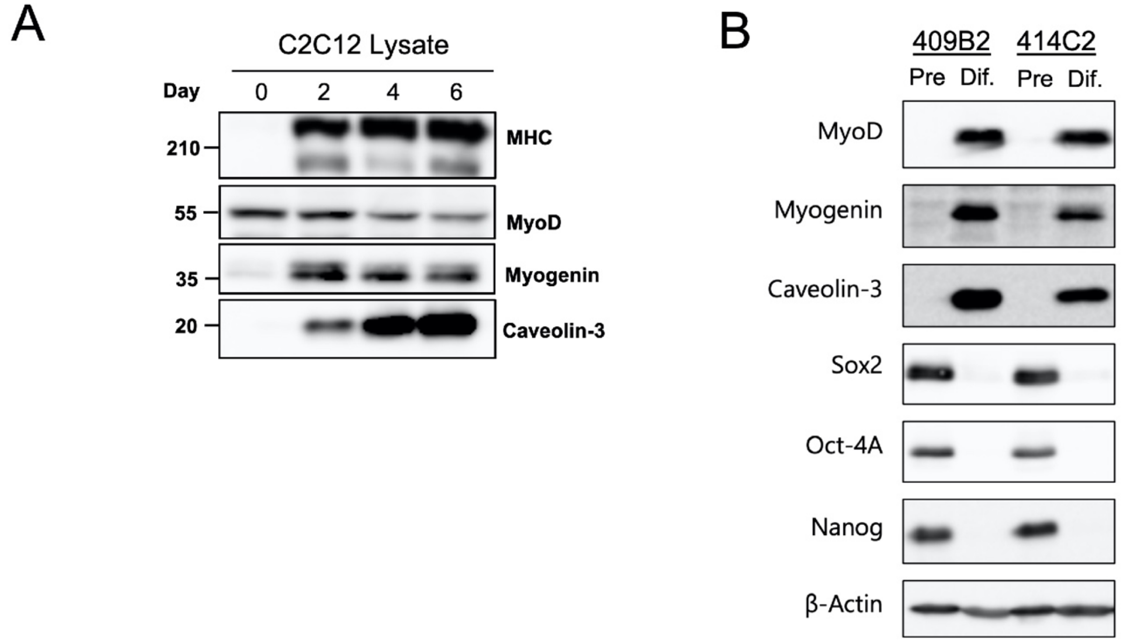


**Fig. S1. Differentiation of C2C12 myoblasts and hiPSCs**

(**A**) Expression of skeletal muscle marker proteins during differentiation of C2C12 myoblasts. Cell lysates were prepared on the indicated time points and subjected to immunoblot analysis. (**B**) Expression of pluripotent and myocyte marker proteins in hiPSC409B2$^{tet\text{-}MyoD}$ and hiPSC414C2$^{tet\text{-}MyoD}$. Cell lysate was prepared on day 0 (pre) and day 5 (Dif), and expression of indicated proteins was analyzed by immunoblot.

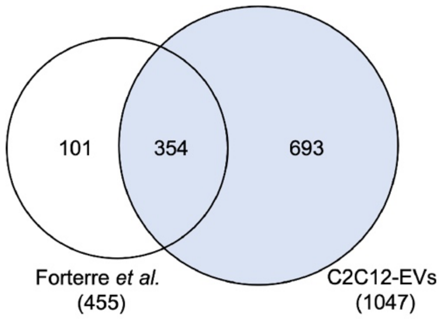


**Fig. S2. Comparison of proteomic analysis on C2C12-derived EVs between current and previous results**

Venn diagram showing overlapping EV proteins from C2C12 cells identified in this study (C2C12-EVs) and the previous study by Forterre et al. (1). Each proteomic data set is derived from both C2C12 myoblast- and myotube-derived EVs.

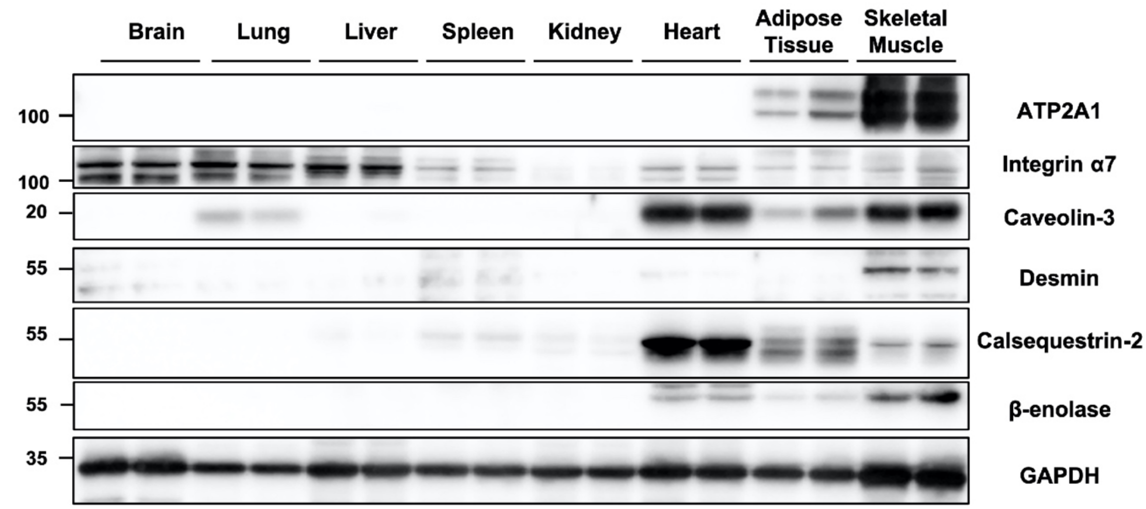


**Fig. S3. Expression of SkM-EV marker proteins in various mouse tissues**

Tissue homogenates were prepared as described in Materials and Methods. Equal amounts of proteins (10 μg/lane) were subjected to immunoblot analysis using the indicated antibodies. GAPDH was detected as a loading control.

A

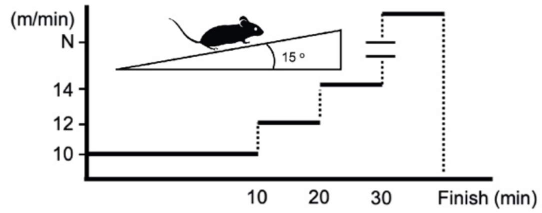


B

| Mouse | Body weight (g) | Run Time (min) |
|---|---|---|
| Ctrl | 22.7 ± 1.0 | - |
| Exercise | 21.3 ± 0.9 | 77.5 ± 11.6 |

C

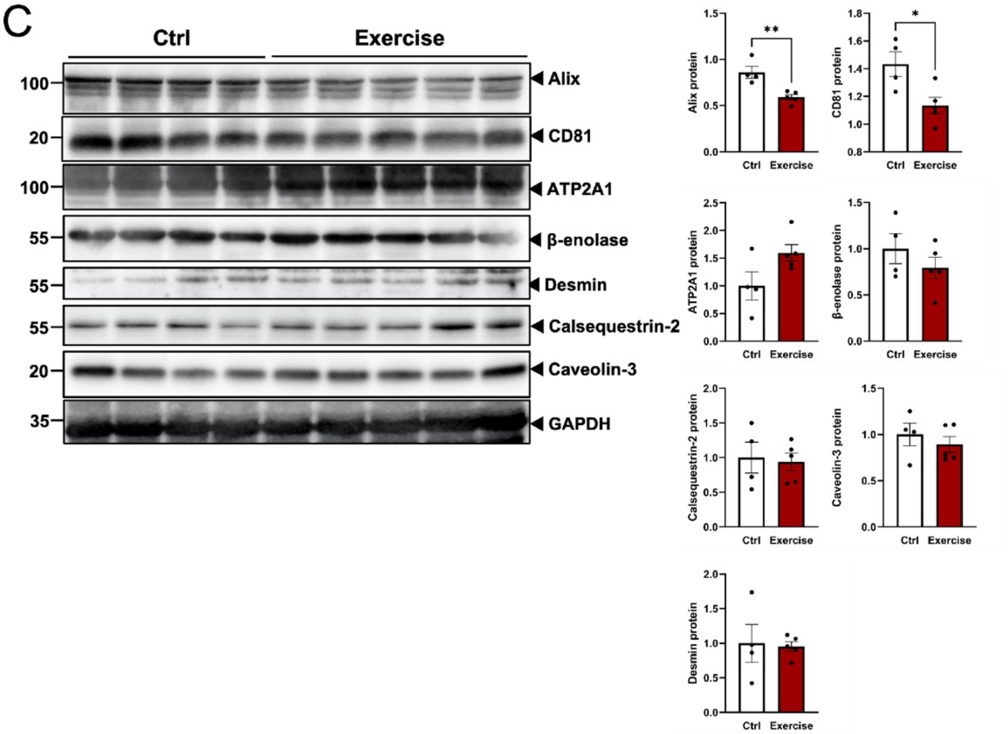


**Fig. S4. Effect of exercise on SkM-EV marker proteins in the skeletal muscle**

(**A**) Schematic presentation of the treadmill running protocol. (**B**) Running time. Mice were subjected to treadmill running as in (A). Results are means ± SD. n=4 for control group, n=5 for exercise group. (**C**) Expression of SkM-EV marker proteins in the skeletal muscle tissue (gastrocnemius) with or without exercise. Left panel: Tissue homogenates were prepared from control and exercised mouse gastrocnemius. Equal amounts of proteins (10 μg) were subjected to immunoblotting. Right panel: Quantification of the protein expression. Protein expression levels were normalized to GAPDH. Results are shown as means ± SEM. Each dot represents individual mice. *p < 0.05, **p < 0.01, n = 4 for control group, n = 5 for exercise group. See Figure 5 for more detail.

**Table S1. The thirty-seven proteins detected in EVs from C2C12 myotubes and hiPSC-myocytes but not from C2C12 myoblasts.**

| UniProt Accession# | | Gene name | Descripition |
|---|---|---|---|
| Mouse | Human | | |
| Q08943 | Q08945 | SSRP1 | FACT complex subunit |
| A2AQA9, A2AQB2 | H0Y786 | NEB | Nebulin |
| Q8BFY9 | Q92973 | TNPO1 | Transportin-1 |
| P50518 | P36543 | ATP6V1E1 | V-type proton ATPase subunit E 1 |
| P62751 | P62829 | RPL23 | 60S ribosomal protein L23 |
| Q91XL3 | Q8NBZ7 | UXS1 | UDP-glucuronic acid decarboxylase 1 |
| A2AUC9 | O60662 | KLHL41 | Kelch-like protein 41 |
| Q5SX40 | P12882 | MYH1 | Myosin-1 |
| Q3TVI8 | Q96AQ6 | PBXIP1 | Pre-B-cell leukemia transcription factor-interacting protein 1 |
| Q91YP3 | Q9Y315 | DERA | Deoxyribose-phosphate aldolase |
| Q02789 | Q13698 | CACNA1S | Voltage-dependent L-type calcium channel subunit alpha-1S |
| E9PYH2 | O00154 | ACOT7 | Cytosolic acyl coenzyme A thioester hydrolase |
| Q9JMA1 | P54578 | USP14 | Ubiquitin carboxyl-terminal hydrolase 14 |
| Q8VHM5 | O43390 | HNRNPR | Heterogeneous nuclear ribonucleoprotein R |
| Q7TMM9 | Q13885 | TUBB2A | Tubulin beta-2A chain |
| O88477 | Q9NZI8 | IGF2BP1 | Insulin-like growth factor 2 mRNA-binding protein 1 |
| Q9Z1E4 | P13807 | GYS1 | Glycogen [starch] synthase, muscle |
| F8VPN4 | P35573 | AGL | Glycogen debranching enzyme |
| Q6PDG5 | F8VXC8 | SMARCC2 | SWI/SNF complex subunit SMARCC2 |
| P29788 | P04004 | VTN | Vitronectin |
| Q3UZG4 | Q12904 | AIMP1 | Aminoacyl tRNA synthase complex-interacting multifunctional protein 1 |
| Q3UQ28 | Q92626 | PXDN | Peroxidasin homolog |
| O88809 | A8K340 | DCX | Neuronal migration protein doublecortin |
| Q9ESE1 | P50851 | LRBA | Lipopolysaccharide-responsive and beige-like anchor protein |
| Q99L88 | Q13884 | SNTB1 | Beta-1-syntrophin |
| Q6P5F9 | O14980 | XPO1 | Exportin-1 |
| Q8BU30 | P41252 | IARS1 | Isoleucine--tRNA ligase |
| P97313 | P78527 | PRKDC | DNA-dependent protein kinase catalytic subunit |
| Q1XH17 | Q6ZMU5 | TRIM72 | Tripartite motif-containing protein 72 |
| Q9D6F9 | P04350 | TUBB4A | Tubulin beta-4A chain |
| Q8BHN3 | Q14697 | GANAB | Neutral alpha-glucosidase AB |
| P70399 | Q12888 | TP53BP1 | TP53-binding protein 1 |

| | | | |
|---|---|---|---|
| P68134 | P68133 | ACTA1 | Actin, alpha skeletal muscle |
| E9PZD8 | P00450 | CP | Ceruloplasmin |
| F8WJ93 | B5MBZ0 | EML4 | Echinoderm microtubule-associated protein-like 4 |
| P10630 | Q14240 | EIF4A2 | Eukaryotic initiation factor 4A-II |
| P70402 | Q13203 | MYBPH | Myosin-binding protein H |

**Table S2. The thirty-one proteins detected only in C2C12 myoblast-derived EVs**

| Uniprot Accession# Mouse | Gene name | Description |
|---|---|---|
| Q8VDZ4 | ZDHHC5 | Palmitoyltransferase ZDHHC5 |
| Q99JR5 | TINAGL1 | Tubulointerstitial nephritis antigen-like |
| P23242 | GJA1 | Gap junction alpha-1 protein |
| Q9WVL3 | SLC12A7 | Solute carrier family 12 member 7 |
| Q9QZM4 | TNFRSF10B | Tumor necrosis factor receptor superfamily member 10B |
| Q60932 | VDAC1 | Voltage-dependent anion-selective channel protein 1 |
| P35700 | PRDX1 | Peroxiredoxin-1 |
| Q8R0W6 | NDFIP1 | NEDD4 family-interacting protein 1 |
| Q8CFE6 | SLC38A2 | Sodium-coupled neutral amino acid transporter 2 |
| Q8C863 | ITCH | E3 ubiquitin-protein ligase Itchy |
| P35288 | RAB23 | Ras-related protein Rab-23 |
| Q80U72 | SCRIB | Protein scribble homolog |
| Q922U2 | KRT5 | Keratin, type II cytoskeletal 5 |
| Q61704 | ITIH3 | Inter-alpha-trypsin inhibitor heavy chain H3 |
| Q9DC51 | GNAI3 | Guanine nucleotide-binding protein G(k) subunit alpha |
| O88952 | LIN7C | Protein lin-7 homolog C |
| O35379 | ABCC1 | Multidrug resistance-associated protein 1 |
| Q8K1S3 | UNC5B | Netrin receptor UNC5B |
| B1ASP2 | JAK1 | Tyrosine-protein kinase |
| Q9D7M5 | DYNAP | Dynactin-associated protein |
| Q61781 | KRT14 | Keratin, type I cytoskeletal 14 |
| O88693 | UGCG | Ceramide glucosyltransferase |
| E9PZW8 | MYO9B | Unconventional myosin-IXb |
| O54890 | ITGB3 | Integrin beta-3 |
| Q9JHF5 | TCIRG1 | V-type proton ATPase subunit a |
| Q8BPM0 | DAAM1 | Disheveled-associated activator of morphogenesis 1 |
| P52293 | KPNA2 | Importin subunit alpha-1 |
| Q9DBH0 | WWP2 | NEDD4-like E3 ubiquitin-protein ligase WWP2 |
| P20934 | EVI2A | Protein EVI2A |
| P58242 | SMPDL3B | Acid sphingomyelinase-like phosphodiesterase 3b |
| Q6IFX2 | KRT42 | Keratin, type I cytoskeletal 42 |

**Table S3. Correlation between EV marker proteins and SkM-EV marker proteins for the interstitium EVs.**

| SkM marker vs EV marker | β-enolase | ATP2A1 | Calseq-2 | Desmin | Cav3 |
|---|---|---|---|---|---|
| Alix | 0.193 | 0.266 | -0.036 | 0.153 | 0.017 |
| CD81 | 0.754 | 0.449 | 0.238 | 0.097 | 0.679 |

The Pearson correlation coefficient was calculated based on Figure 5a.

**Table S4. Primer lists.**

| Primer | Sequence (from 5' to 3') |
|---|---|
| 18S rRNA_Forward | ACCGCAGCTAGGAATAATGGA |
| 18S rRNA_Reverse | GCCTCAGTTCCGAAAACCA |
| Cyclin B1_Forward | CAGAGTTCTGAACTTCAGCCTG |
| Cyclin B1_Reverse | TTGTGAGGCCACAGTTCACCAT |
| Cyclin D1_Forward | GCCGAGAAGTTGTGCATCTACA |
| Cyclin D1_Reverse | TGTTCACCAGAAGCAGTTCCATT |
| Myh1_Forward | CCAAGGGCCTGAATGAGGAG |
| Myh1_Reverse | GCAAAGGCTCCAGGTCTGAG |
| MyoD1_Forward | GCTTCTATCGCCGCCACTCC |
| MyoD1_Reverse | CGCACATGCTCATCCTCACG |
| Myogenin_Forward | GCATGTAAGGTGTGTAAGAG |
| Myogenin_Reverse | GCGCAGGATCTCCACTTTAG |
| Myf5_Forward | GATGTGGGCCTGCAAAGC |
| Myf5_Reverse | TGCGCCGATCCATGGTA |
| Pax7_Forward | TCCCGTCAGCTCCGTGTT |
| Pax7_Reverse | TCCTGATATCGGCACAGAATCTT |
